# Temporal stasis in three facets of biodiversity reveals mechanisms of community assembly across elevation gradients

**DOI:** 10.64898/2026.10.05.756846

**Authors:** Kaleb A. Goff, Michael J. Koontz, Catherine Laufenberg, Sophie Meng, Malia Reiss, Jan Nachlinger, Catie Bishop, Jim Bishop, Mary T. Burke, Brian V. Smithers, Meagan F. Oldfather, Seema N. Sheth

## Abstract

Understanding spatial and temporal variation in community composition is central to protecting and monitoring biodiversity. Similarity in the composition of two communities decreases along spatial and temporal gradients, a relationship known as distance decay. Comparing distance-decay relationships across taxonomic, functional, and bioclimatic facets of diversity provides a means of interrogating niche-based versus neutral influences on community assembly. Furthermore, the context in which these facets yield overlapping or complementary information about global change impacts remains poorly understood. In this study, we examined spatiotemporal distance-decay relationships for taxonomic, functional, and bioclimatic facets of alpine plant diversity leveraging elevation gradients on five peaks in the White Mountains, California, USA, sampled at 3-4 time points between 2007 and 2024. Using multivariate Bayesian beta regression models, we estimated the slopes of elevation distance decay over multiple surveys and across taxonomic, functional and bioclimatic diversity to understand how spatial distance-decay relationships are changing through time and across different facets of diversity. We found strong spatial distance decay of taxonomic similarity across all peaks, with the highest elevation peak showing the strongest decay. Functional and bioclimatic similarities decayed more weakly than taxonomic similarity, suggesting environmental filtering shapes trait and niche distributions. Critically, spatial decay patterns showed no temporal signal across facets of diversity, indicating community stasis rather than biotic homogenization or differentiation. This study adds further evidence that xeric alpine plant communities are resisting climate change impacts through time.

## Introduction

The rate at which similarity varies along spatial and temporal gradients explains how the environment, dispersal, and stochasticity interact to shape communities (Blowes *et al*. 2019; Whittaker 1960). A ubiquitous observation in ecology is that communities become less similar as spatial or temporal distance between communities increases–a pattern known as distance decay (Nekola & White 1999). Distance-decay relationships are the product of species turnover which is driven by underlying community assembly mechanisms, including niche-based species sorting by the environment, dispersal barriers in the landscape, and ecological drift (Anderson *et al*. 2011; Dornelas *et al*. 2014; Heino *et al*. 2024; Qian *et al*. 2005). Distance-decay functions are applicable to a variety of temporal and spatial gradients, evaluating how community similarity changes across spatial scale (Soininen *et al*. 2007), latitude (Bahram *et al*. 2013), elevation (Wang *et al*. 2022), along various environmental gradients (Tuomisto *et al*. 2003), and across time (Collins *et al*. 2000). Importantly, comparing spatial and temporal distance-decay patterns simultaneously reveals additional information. For example, examining spatial decay relationships across multiple time points can reveal whether drivers such as a changing climate are restructuring spatial patterns through time; while examining temporal decay across multiple spatial gradients can explain what places are changing more or less rapidly (Dallas *et al*. 2024).

The rate of change of spatial distance-decay relationships across multiple time points can be used to understand community trajectories (Bagchi & others 2017; Collins *et al*. 2000; Rolls *et al*. 2023). Mechanisms that reduce environmental heterogeneity across a landscape, such as climate warming, can cause species turnover that results in the convergence of community composition over time, weakening spatial distance-decay slopes in a pattern termed biotic homogenization. (Chen et al. 2025; Magurran et al. 2015, Figure 1). Conversely, if local environmental filtering strengthens over time, spatial distance-decay slopes become steeper, a pattern termed biotic differentiation (Olden & Rooney 2006, Figure 1). Finally, if dispersal, colonization and extinction lags are long, communities shift synchronously, or if there is strong topographic buffering, spatial distance-decay relationships may remain stable across surveys despite ongoing environmental change, consistent with stasis (Figure 1). Thus, evaluating temporal signals in spatial decay is a powerful way to inform biodiversity monitoring in the wake of accelerating climate change, but to date has rarely been quantified.

**Figure 1:**
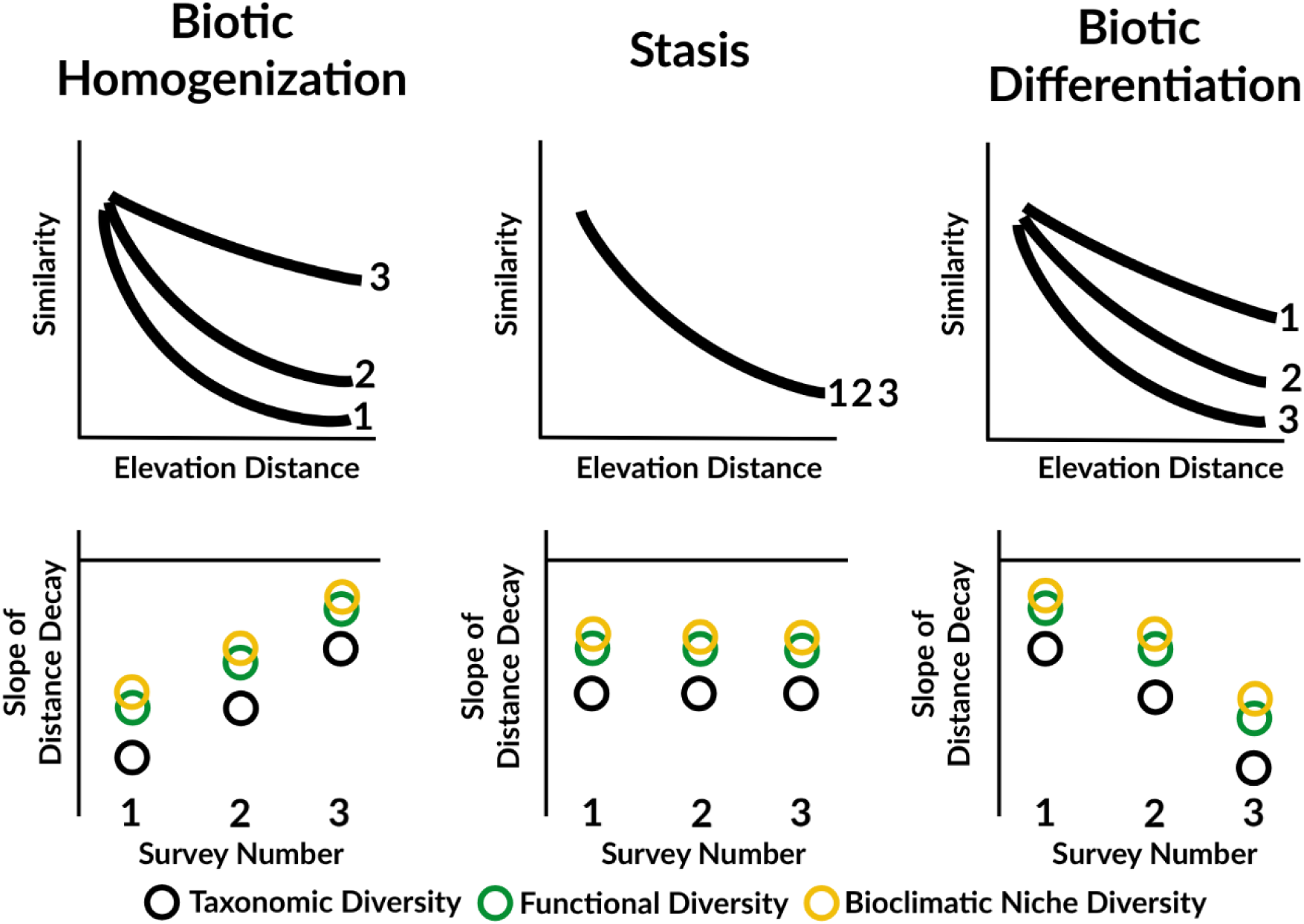
Conceptual diagram showing hypotheses for the rate of distance decay across three facets of diversity: taxonomic, functional and bioclimatic niche (**H2**), and temporal signals in elevation distance decay (**H3**). We hypothesize that functional and climatic niche similarities show a weaker elevation distance decay than species composition, because in high elevation systems extreme abiotic conditions drive functional replacement across elevation gradients via the clustering of species traits (bottom row, distance between black and green and yellow circles). In addition, we hypothesize three contrasting scenarios related to the temporal signal in elevation distance decay driving differences in the rate of community change (**H3**). **Homogenization (H3.1):** A temporal signal in elevation distance decay whereby the slopes of elevation distance-decay relationships on different peaks become shallower over multiple chronological surveys. Ecologically, this is caused by plant communities that are further away from one another along the elevation gradient becoming more similar to one another over time. **Stasis (H3.2):** No temporal signal in elevation distance decay, related ecologically to lags in dispersal, recruitment and/or colonization. **Directional Change (H3.3):** A temporal signal in elevation distance decay where slopes of elevation distance-decay relationships on each peak become steeper over multiple chronological surveys. Ecologically, this is due to greater differences between plant communities along the elevation gradient.

Communities can be described through multiple facets of diversity, such as taxonomic (species identities), functional (species traits), and bioclimatic (species climate niches) — each of which varies in its sensitivity to niche-based versus neutral assembly processes (Jarzyna & Jetz 2016; Cadotte et al. 2011). When niche-based assembly processes govern community assembly along a strong spatial or temporal gradient, differing conditions at each end of the gradient select for distinct functional and bioclimatic strategies, driving faster distance decay in functional and bioclimatic composition than in taxonomic identities (Swenson et al. 2011). Alternatively, if functional and bioclimatic strategies are more conserved across the species pool than species identities, or if the same limited range of strategies are favored along a gradient, niche-based facets may similarly decay more slowly than taxonomic composition (Heino & Tolonen 2017). Further, If environmental filtering is strong, communities may converge in terms of species traits but diverge in terms of species identities (Fukami et al. 2005) Finally, when dispersal limitation drives community turnover independent of species traits, all three facets should decay at similar rates. Comparing distance-decay relationships across facets therefore provides a window into the relative influence of niche-based versus neutral processes in structuring communities across space and time, an approach that remains underutilized in biodiversity monitoring (Graco-Roza et al. 2022).

Evaluating knowledge gaps about distance decay across spatial and temporal gradients and for multiple facets of diversity requires a study system with strong spatial and environmental gradients, undergoing some process of change over time. Elevation gradients are spatial gradients with multiple underlying environmental gradients, and high-elevation systems are experiencing rapid and accelerating warming over time due to climate change (Hock *et al*. 2019). Elevation gradients contain high biodiversity across multiple facets because they compress climatic gradients over short spatial extents, include topographic heterogeneity, and are shaped by historical contingency (IPBES 2019). Impacts to mountain biodiversity from climate change have the potential to result in biotic homogenization (Britton *et al*. 2009; Savage & Vellend 2015), stasis (Goff *et al*. 2025) or biotic differentiation (Gottfried *et al*. 2012; Steinbauer *et al*. 2018). We expect that because environmental conditions become increasingly harsh and selective with increasing elevation, alpine areas will provide clear signals of the impact of environmental filtering and reduced species pool size on distance-decay relationships. The stringent abiotic conditions at high elevations also create strong trait and niche clustering, providing a test of whether functional and bioclimatic similarities decay slower than taxonomic composition with increasing elevation distance.

We leverage a spatiotemporal dataset of elevation gradients on five peaks in the White Mountains, California, USA), to address the following hypotheses. First, we hypothesize that higher elevation peaks have steeper elevation distance-decay relationships, due to fewer overall species and stronger environmental filtering (**H1**). Second, we hypothesize that functional and climatic niche similarities show a weaker elevation distance decay than species composition, because in high elevation systems, extreme abiotic conditions drive clustering of species traits and bioclimatic niches (**H2**, Figure 1). Third, we hypothesize a temporal signal in these elevation distance-decay relationships will conform to one of three contrasting scenarios: biotic homogenization (**H3.1**), stasis (**H3.2**), and biotic differentiation (**H3.3**, Figure 1).

## Materials and Methods

### Study Area and Plant Community Data

A detailed description of the study area, and the field methods for the plant community data can be found in (Smithers and Oldfather *et al*. 2020). Briefly, plant community data for this study were collected following on-contour belt transects situated along elevation gradients included as an extra approach in the GLORIA (Global Observation Research Initiative in Alpine Environments) field manual (Pauli *et al*. 2015). On five peaks in the White Mountains, California, USA, 100 m x 1 m belt transects were established in 2007 (SME, Sheep Mountain East), 2008 (WMT, White Mountain Peak and PGS, Patriarch Grove South), and 2011 (CPT, Campito Mountain and BAR, Mount Barcroft), starting at 35 vertical meters downslope of the highest summit point and continuing downslope at intervals of 25 vertical meters (Figure 2). The total length per peak of these downslope elevation gradients ranges from 200-350 vertical meters. Surveys of the downslope transects were completed at multiple time points at approximately 5-year intervals, with a total of 3-4 surveys for each peak for a total time span of 2007-2024. Each 100 m belt transect was divided into ten 10 m x 1 m segments, and within each segment, we recorded all plant species. We use the total number of segments per transect where the species occurred as a measure of species abundance (Figure 2).

**Figure 2:**
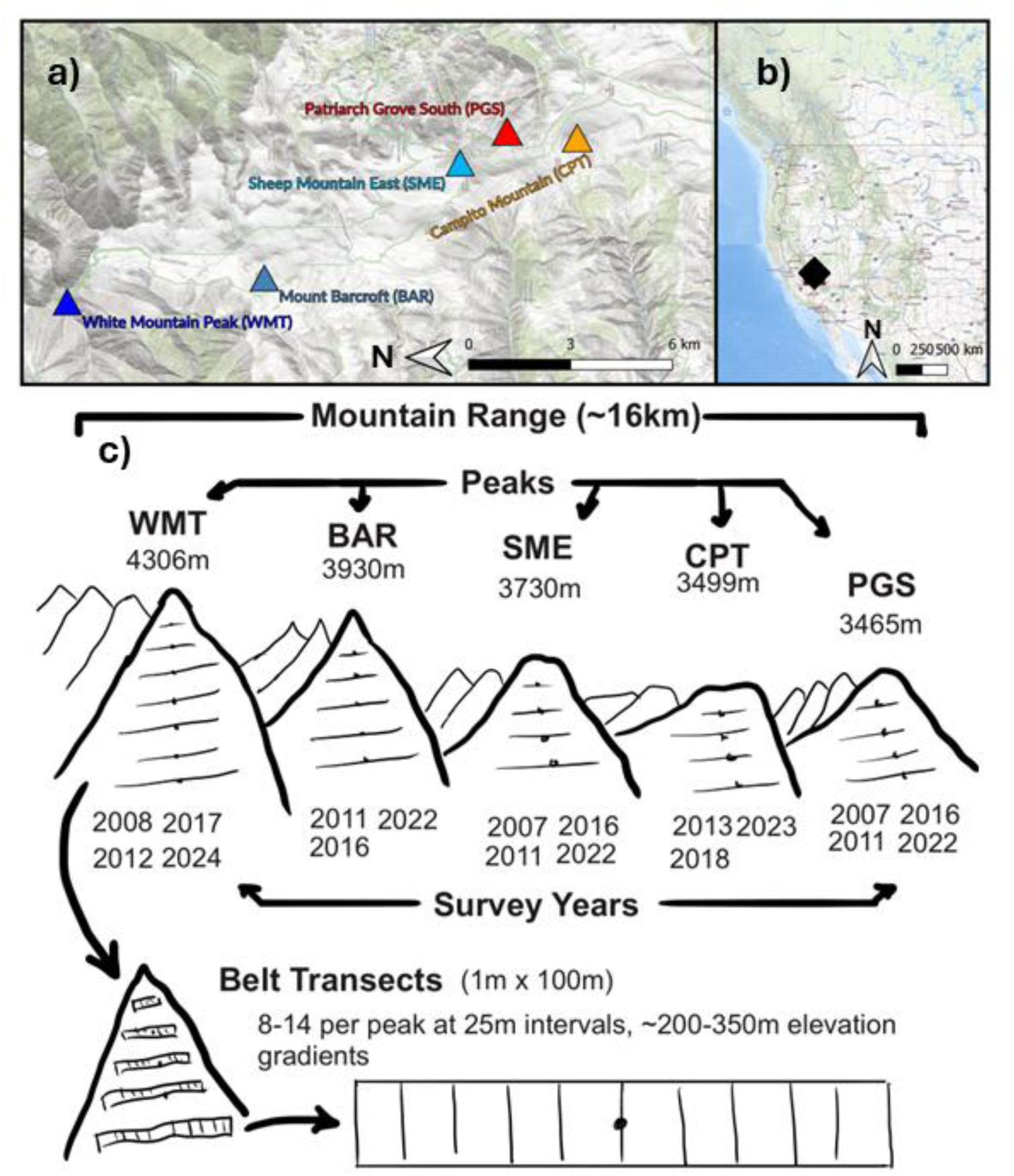
Study area map: **(a)** peaks relative to one another and **(b)** study area in western North America) and study design schematic **(c)**. On five peaks in the White Mountains, CA, USA, plant community data were collected in 100 m x 1 m belt transects placed on contour at 25m intervals. These transects on the peaks were sampled across multiple survey years. Within each transect, we collected data on the presence/absence of all species in ten 10 m x 1 m segments. We estimated the abundance of each species on each transect in each year as the number of segments on which the species was present.

### Functional Trait and Bioclimatic Niche Data

We conducted field campaigns in the White Mountains to assemble a dataset of locally collected functional traits related to the plant economic spectrum (Reich 2014) (details in Supporting Information, Appendix S1). Briefly, we collected data on maximum plant vegetative height (cm), leaf wet mass (g), leaf dry mass (g) and leaf area (cm^2^), specific leaf area (cm/g) and leaf dry matter content (g/g) using standardized methods (Perez-Harguindeguy *et al*. 2016). We assembled mean trait values for 92 species representing 95.3% of the total plant abundance across all transects and years. For each species, we collected measurements of each trait on at least nine individual plants, sampling across each species’ elevation range within the White Mountains. Trait data collection took place in 2022, 2023 and 2024. We included only maximum plant vegetative height (cm), specific leaf area (cm/g) and leaf dry matter content (g/g) in the analyses, to reduce collinearity between traits.

To describe the bioclimatic niche of each species in the community data, we created an occurrence dataset from each species’ North American distribution using data from the Global Biodiversity Information Facility (GBIF.org 2024) and combined these occurrence points with climate data from CHELSA (Climatologies at High resolution for the Earth’s Land Surface Areas; (Brun *et al*. 2022). First, we downloaded all vouchered specimen records from the North American distribution of each species in the community data using the R package “rgbif” (Chamberlain *et al*. 2024). These occurrences were cleaned using automated methods in the R package “coordinatecleaner” (Zizka *et al*. 2019), and then manually cleaned by mapping occurrence points in R and investigating outlying points. Next, we downloaded gridded climate norm data from CHELSA at 30 arc-second resolution representing mean values across the 1981-2010 period, and calculated spatial summaries of the mean, median and range of 10 bioclimatic variables related to averages, variability, and extremes, chosen based on our knowledge of the system, across the occurrences of each species (Fick & Hijmans 2017; Karger *et al*. 2017). To reduce collinearity between predictors we removed climate variables with correlations greater than 0.75, reducing from 10 bioclimatic variables to 5 (mean annual temperature (°C), mean annual precipitation (mm), isothermality (mean of monthly max - min), temperature seasonality (°C), and precipitation seasonality (mm). A detailed description of these methods can be found in Supporting Information, Appendix S2.

### Taxonomic Harmonization Across Facets of Diversity

We standardized species names, removed intraspecific ranks and aggregated problematic species complexes across the three datasets to reduce pseudoturnover (Boch *et al*. 2022). Specifically, we cleaned and harmonized taxon names using the White Mountains Vascular Plant Checklist (Morefield *et al*. 2022). To match the three datasets (plant community, functional traits and species bioclimatic niche) and ensure we queried the maximum number of occurrences from GBIF, we used binomial species names throughout. Across all three datasets, we aggregated species complexes that were difficult to identify over the study period into “dummy” species (Grenié *et al*. 2023; Zhu *et al*. 2024). A detailed description of these methods, including specific dummy species aggregations, can be found in Supporting Information, Appendix S3. At the end of the harmonization process, we had a total of 130 species in the community data, bioclimatic niche means for 129 species, and functional traits for 92 species.

### Statistical Analyses

We performed all data analyses in R (R Core Team 2025) and made use of the R packages “tidyverse” (v2.0.0;(Wickham *et al*. 2019) and “stats” (v4.4.0; R Core Team, 2025). We used a Bayesian statistical modeling framework, running models in the R package “brms” (v2.23.0; (Bürkner 2021), and using the R package “marginaleffects” for computing marginal predictions and slopes (v0.31.0; (Arel-Bundock *et al*. 2024).

To measure the rate of decay of taxonomic similarity, we first calculated pairwise Sørensen dissimilarity including species abundances using the function *beta* in the package “BAT” (v2.9.6, (Cardoso *et al*. 2024) and subtracted 1 from each pairwise comparison to convert to similarity, with 1 being completely similar communities, and 0 being completely dissimilar communities. Next, we filtered pairwise distance measures to include only comparisons between transects at the same peak in the same year, excluding self-comparisons. For example, similarity measures for Mount Barcroft include pairwise comparisons between all 14 belt transects along the elevation gradient for 2011, then another set of similarity measures for 2016, and another for 2022. We could thus examine changes in the elevation distance-decay relationship on peaks across multiple time points.

To model the rate of decay of functional and bioclimatic similarity, we first computed the dissimilarity among species across multiple functional traits and bioclimatic niche axes. For functional traits, we calculated multi-trait dissimilarity using mean trait values of maximum plant height (cm), specific leaf area (cm/g) and leaf dry matter content (g/g) across all species, using the R package “gawdis” (v0.1.5; (de Bello & others 2021). We then reduced these multi-trait dissimilarities to three PCoA axes using the function *cmdscale* from the R package “stats”, where 81% of the variance in trait dissimilarity among species was explained by the first three axes. Using the three PCoA axes, we built kernel density hypervolumes of functional trait diversity using the function *kernel.build* from the R package “BAT” (Cardoso *et al*. 2024), including species abundances. We then used the function *kernel.beta* from the R package “BAT” to compare the overlap of hypervolumes, where each hypervolume represents the functional diversity of a belt transect on one peak at one timepoint (see Figure 2 from (Mammola & Cardoso 2020). We subtracted 1 from each measure of hypervolume overlap to convert to similarity and filtered these measurements to include only comparisons between belt transects on the same peak in the same year. We repeated this process to measure the similarity of bioclimatic niches. We then constructed the climatic niche hypervolumes using means for each species of mean annual temperature (°C), mean annual precipitation (mm), isothermality (mean of monthly max - min), temperature seasonality (°C), and precipitation seasonality (mm).

We modeled changes in taxonomic, functional and bioclimatic similarity across elevation gradients and surveys using multivariate beta regression implemented in the R package “brms” (Bürkner 2021). We accomplished this by constructing separate submodels for similarity of each facet of diversity, using the same submodel formulation for each response variable. Fixed effects included the difference in elevation between two belt transects, survey number and their interaction. To account for among-peak variation, each submodel also included random intercepts and random slopes for elevation difference, survey number, and their interaction, for each peak. These three submodels were combined into a single multivariate formulation, allowing submodels to share correlated peak-level random effects while assuming independent residuals across responses. Similarity values did not include exactly zero or one, so a standard beta likelihood was appropriate and zero–one-inflated extensions were unnecessary. We selected priors based on our knowledge of the system and evaluated priors using prior predictive checks. Posterior distributions were sampled using Markov Chain Monte Carlo with four chains, each with 2,000 warm-up iterations followed by 2,000 sampling iterations, using an initial value of 0. Model fitting was performed with the cmdstanr backend (v0.9.0; (Gabry *et al*. 2025), with control parameters set to adapt_delta = 0.998 and max_treedepth = 14 to ensure stable exploration of the posterior and no divergent transitions.

With the posterior distributions from similarity models for taxonomic, functional trait, and bioclimatic facets of diversity, we used the function *predictions* in the R package “marginaleffects” to make model predictions for distance decay curves for each peak, survey number and facet of diversity. From these curves, we calculated the average slope of the elevation distance-decay relationship using the function *avg_slopes*. To address our first hypothesis, we compared elevation distance-decay slope means and 95% credible intervals (CI) among peaks of differing median elevations (H1). Next, we addressed differences in slopes among facets of diversity (H2), by extracting joint posterior draws of the elevation slope and its interaction with survey number for each peak from the multivariate model. Conditional slopes for each survey were calculated by summing population-level fixed effects and peak-level random deviations for both main and interaction terms. Paired differences between facets were computed for each posterior draw, preserving correlations across responses. Where mean posterior differences and 95% credible intervals overlap zero, we conclude that there is no difference in slopes between two facets of diversity. If the difference is negative, the first facet in the comparison has a slope that is more negative/steeper than the second facet in the comparison, and if the difference is positive, the first facet in the comparison has a less negative or shallower slope than the second facet. Lastly, to evaluate hypothesis 3, we obtained the interaction between survey number and elevation distance by summing the population-level interaction term (fixed effect) with the peak-level deviations for the interaction effect for each peak (random effects). Positive interaction effects suggest a dampening of the negative distance-decay relationship, consistent with biotic homogenization (H3.1). Negative interaction effects are consistent with biotic differentiation (H3.3), whereby negative distance-decay slopes are becoming steeper, and stasis is evidenced by 95% CIs of the interaction effect overlapping zero (H3.2).

## Results

### Strongest spatial distance decay on the highest elevation peak (H1)

The distance decay of species identities across elevation was consistently negative for all peaks (Figure 3a). The average marginal effect of elevation difference across all peaks and years for the distance decay of taxonomic similarity was -0.00162 (95% credible interval: - 0.00172 to -0.00153), or a -0.162% difference in similarity per meter of elevation distance. Transects were placed at 25m intervals, meaning that on average community compositional similarity decreased by -4.05% (95% credible interval: -3.82% to -4.29%) for each step along the elevation gradient (with 8 - 14 steps depending on peak). The highest elevation peak (WMT) had the steepest taxonomic distance-decay relationship (average slope -0.00224, 95% credible intervals: -0.00211 to -0.00237), while three of the four other peaks had smaller slope values that overlapped one another (BAR, PGS and SME, Table 1). The second lowest elevation peak, CPT, had the widest credible intervals for the effect of elevation difference, and an effect between the middle and highest peaks (-0.00167, 95% credible interval: -0.00121 to -0.0021). The spatial distance decay of functional traits was consistently negative across all peaks and years, except on the lowest elevation peak, where credible intervals overlapped zero (Figure 3b, Figure 4a). The average marginal effect of elevation difference in the functional decay model was -0.00092 (95% credible interval: -0.00103 to -0.0008), or a -2.29% (95% credible interval: - 1.99% to -2.58%) decrease in functional trait similarity with each 25m step along the elevation gradient. Distance decay of bioclimatic niche similarity was also negative on all but the lowest elevation peak (PGS), where the elevation distance decay credible intervals overlapped zero (Figure 3c, Table 1). The average rate of bioclimatic niche elevation distance decay was - 0.00107 (95% credible interval: -0.00118 to -0.00097), equating to a -2.68% (95% credible interval: -2.42% to -2.94%) decrease in bioclimatic niche similarity with each 25m step along the elevation gradient.

**Figure 3:**
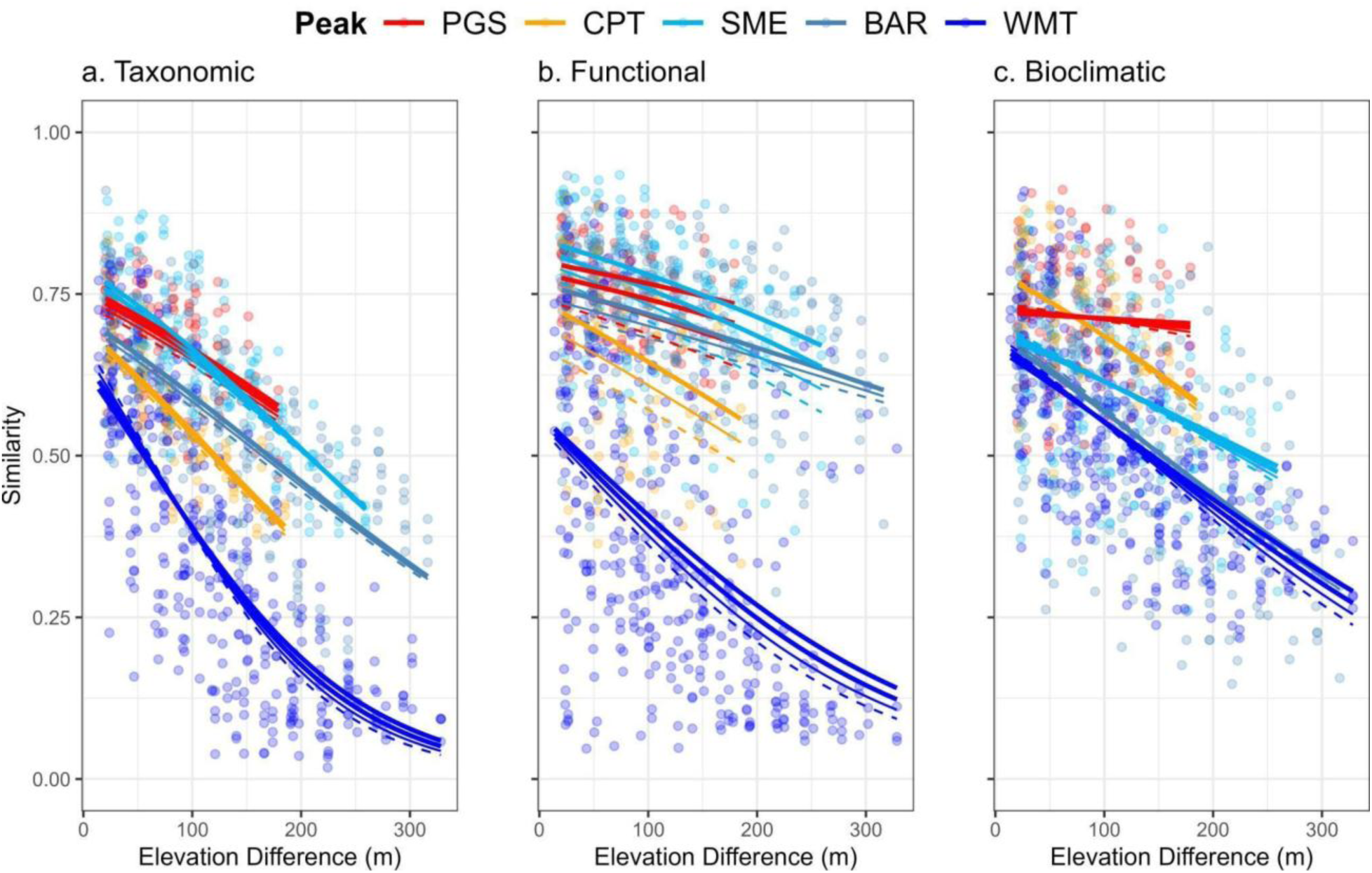
Distance decay curves of taxonomic similarity **(a)**, functional similarity **(b)** and bioclimatic niche similarity **(c)** for multiple peaks (colors) and multiple surveys within each peak (lines) across increasing elevation distance between two transects on the same peak in the same year. Dashed lines represent the 1st surveys of each peak, thin solid lines show the 2nd survey, and thick solid lines represent the 3rd and 4th surveys of each peak.

**Figure 4:**
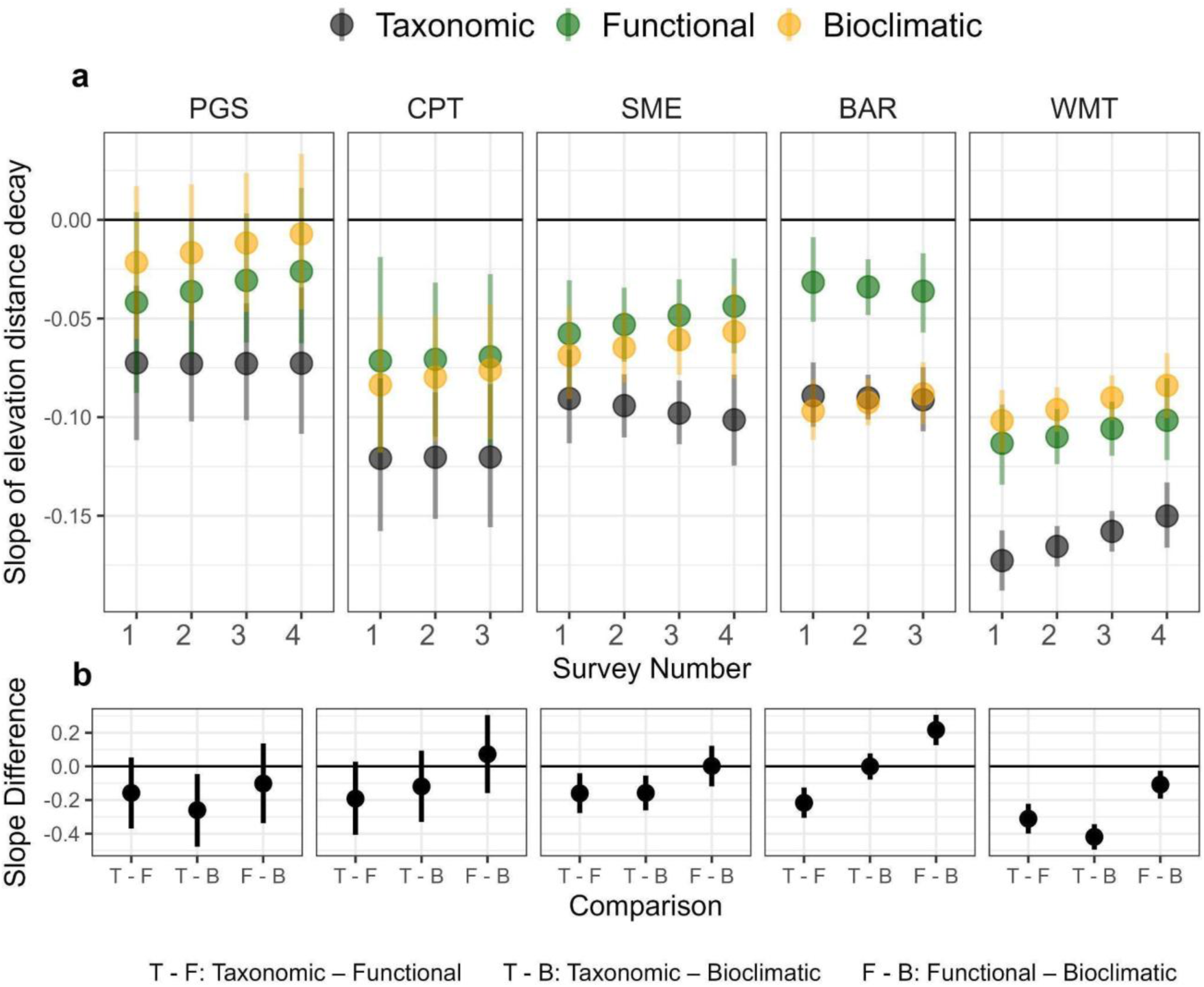
**(a)** Slopes of elevation distance-decay relationships across three facets of diversity: taxonomic (species identities), functional (functional diversity in vegetative height, SLA and LDMC) and bioclimatic niche (suite of climate variables representing averages, variability and extremes), and 3-4 surveys at 5-year intervals of each peak. Line ranges show 95% credible intervals. Facets are arranged by median peak elevation, with the lowest peak on the left and the highest peak on the right. **(b)** Posterior differences in elevation distance-decay slopes between biodiversity facets. Positive values indicate shallower slopes for the first facet; negative values indicate steeper slopes. Points show posterior means; lines show 95% credible intervals.

**Table 1:**
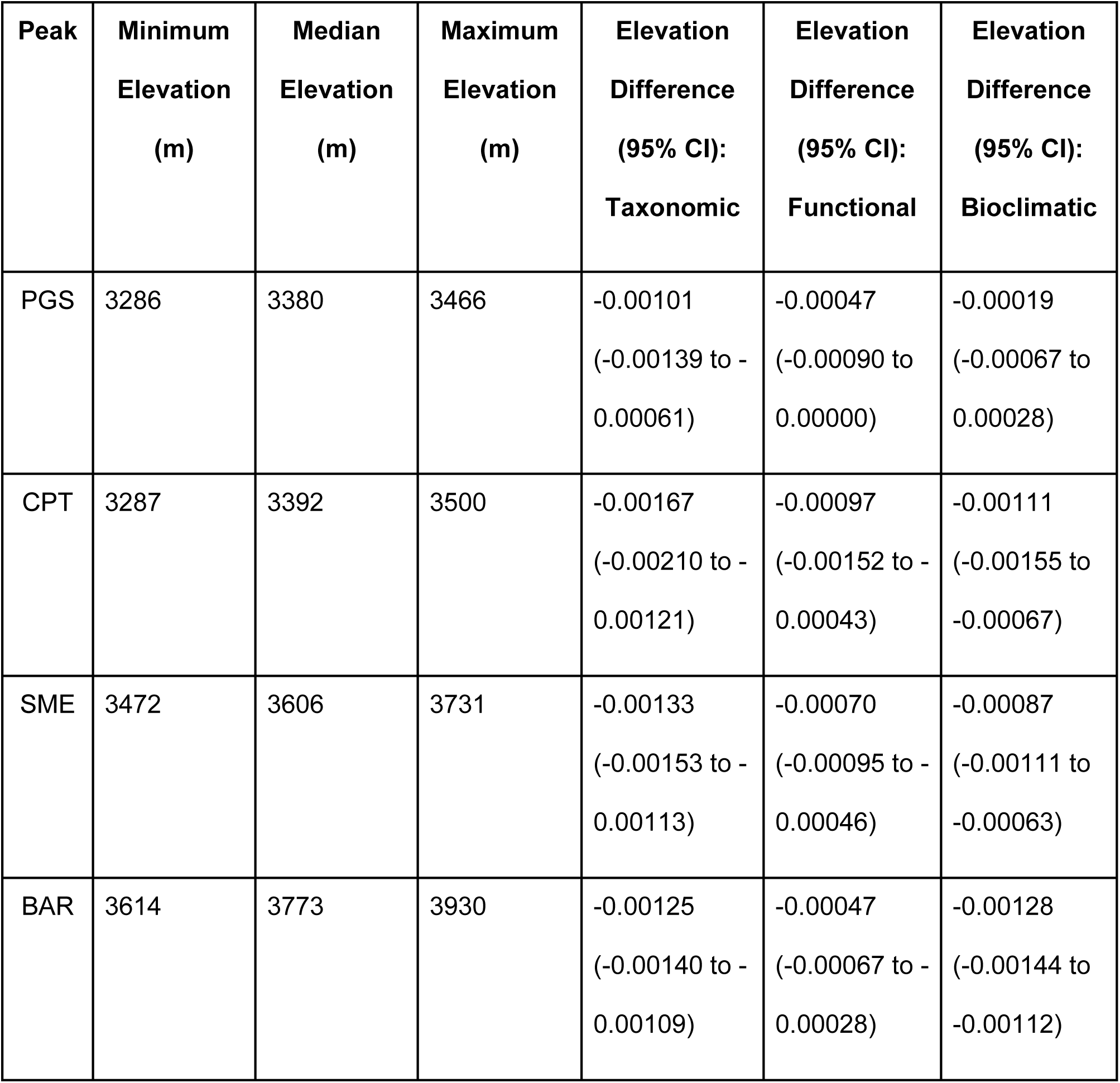
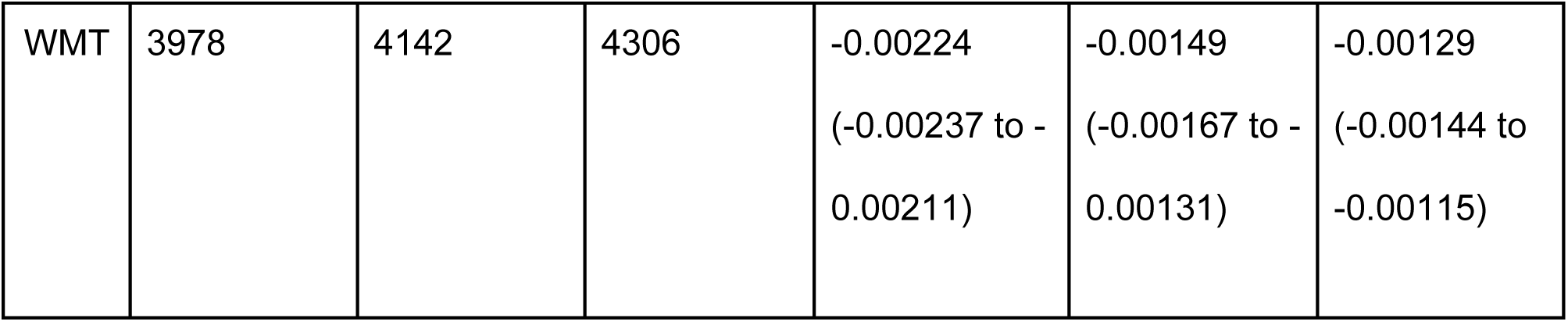
Peak median and max elevation, and results from Bayesian hierarchical model of similarity for three facets of diversity (taxonomic, functional trait and bioclimatic niche), showing the average effect of elevation difference, conditional on each peak, in units of percent change in similarity for each meter of elevation change

### Distance decay of functional similarity is weaker than taxonomic and bioclimatic similarity, but only on the highest elevation peaks (H2)

Functional, bioclimatic and taxonomic distance decay rates differed only on the two highest elevation peaks. On the second highest peak, Mt. Barcroft (BAR), functional trait similarity decayed more slowly than taxonomic similarity (-0.21672, 95% credible interval: - 0.12614 to -0.30601) and bioclimatic niche similarity (0.21676, 95% credible interval: 0.30629 to 0.12624) (Figure 4b). On the highest elevation peak, White Mountain Peak (WMT), taxonomic similarity decayed faster than functional (-0.31114, 95% credible interval: -0.22254 to -0.39938) or bioclimatic similarity (-0.41914, 95% credible interval: -0.34315 to -0.49512), while functional and bioclimatic decay were weaker and overlapping (-0.108, 95% credible interval: -0.02603 to - 0.19138) (Figure 4b). On both peaks, functional decay was weaker than taxonomic decay along elevation gradients.

### Limited temporal signal in spatial distance decay (H3)

We did not detect a temporal signal in elevation distance decay; across all three facets of diversity, 95% credible intervals for the interaction between elevation distance and survey number overlapped one another (Figure 4). The average marginal effect of survey number on taxonomic distance decay across the surveys overlapped zero. (95% credible interval: -0.00046 to 0.01218). The interaction between survey number and elevation difference was weak, with an average taxonomic elevation distance decay slope of -0.00162 (95% credible interval: -0.00177 to -0.00147) in the first survey, and -0.00159 (95% credible interval: -0.0017 to -0.00147) in the third survey. For functional traits and bioclimatic niches, the average marginal effect of survey number overlapped zero (functional traits 95% credible interval: -0.01217 to 0.02823, bioclimatic niche 95% credible interval: -0.00673 to 0.00751). For bioclimatic niches and functional traits, there was no interaction between survey number and elevation distance decay slopes. Finally, the 95% credible interval for the total interaction effect of survey number and elevation distance for each peak overlapped zero for all peaks and facets of diversity (Figure S3).

## Discussion

The goal of this study was to leverage distance-decay relationships to examine spatiotemporal patterns of community change across multiple facets of biodiversity, and relate these patterns to community assembly mechanisms. Our study of distance decay along elevation gradients on five xeric alpine peaks surveyed 3-4 times over 17 years in the White Mountains, CA, USA found the strongest spatial distance-decay relationships for taxonomic similarity. In agreement with our first hypothesis, the highest elevation peak (White Mountain Peak) had the steepest elevation distance decay compared to lower elevation peaks, but only for the taxonomic facet of diversity (Figure 3). We found mixed support for our second hypothesis, that functional and bioclimatic facets of diversity decay slower than species identities, which was observed on two of the five peaks (Figure 4b). Finally, we found evidence for community stasis across the three facets of diversity, rather than biotic homogenization or differentiation across these gradients, even after 3-4 surveys spanning nearly two decades. The distance-decay analysis framework allowed us to examine changes in multiple facets of biodiversity across spatial and temporal gradients simultaneously, and relate these to niche-based vs neutral assembly processes, a major challenge in community ecology. Below, we discuss distance decay comparisons across facets of diversity, the lack of temporal signal in spatial decay, and the distance decay analysis framework from the perspective of global change.

Larger spatial gradients and larger species pools have been shown to produce stronger distance-decay relationships in a variety of habitats and taxonomic groups (see Figure 5 in (Graco-Roza *et al*. 2022). We consistently detected negative distance-decay relationships along 200-350 m elevation gradients, across all three facets of diversity. This supports the assertion that spatial distance decay manifests over relatively short spatial scales. In our study, the highest elevation peak (White Mountain Peak) had the fewest species and the steepest elevation distance-decay relationship. White Mountain Peak and Mount Barcroft represent the longest elevation gradients (350m), but Mount Barcroft includes more species and had a weaker distance-decay relationship, suggesting that in this system, the size of the species pool is more important than differences in elevation gradient length in the rate of spatial distance decay. Finally, larger spatial gradients have stronger decay relationships in part because they cross habitat boundaries (Nekola & White 1999). In this system, White Mountain Peak includes the most dramatic habitat boundary, transitioning from alpine grassland and cushion plants to talus and bare rock, highlighting the importance of extreme environmental conditions and stark habitat boundaries in driving decay of community similarity.

Understanding when and where different facets of biodiversity yield overlapping, complementary, or contrasting information about community change is still a pressing question in community ecology (Heino & Tolonen 2017; Perez Rocha *et al*. 2019). We expected that taxonomic diversity would decay faster than functional and bioclimatic niche diversity because niche-based facets reflect adaptations to environmental constraints that are likely to be shared across species in similar habitats (Fukami *et al*. 2005; Mouillot *et al*. 2013). Taxonomic distance-decay slopes were greater than functional and bioclimatic only on the highest elevation peaks, although there was a trend toward taxonomic decay being steepest on the other peaks (Figure 4b). One likely explanation for this finding is that the communities included in this study are subject to similar environmental filtering and are subsamples of a larger species pool (Sokol *et al*. 2011). In xeric alpine plant communities, habitat constraints are similar, and the spatial scale analyzed is small, so dispersal and species pool effects are stronger for species diversity than functional diversity (Soininen *et al*. 2016). This is consistent with functional and bioclimatic strategies being more conserved across the regional species pool than species identities. In other words, our elevation gradients captured more of the functional and bioclimatic niche space of alpine plants in this study system than the total number of species. Our results are consistent with a recent study that found taxonomic decay to be stronger than functional decay across spatial gradients (Graco-Roza *et al*. 2022). To our knowledge, this is the first analysis of distance decay in bioclimatic niche similarity, and decay rates for this facet of diversity aligned closely with functional trait diversity. Future studies comparing distance-decay patterns in traits and climatic niches would help to further examine this relationship.

Investigating temporal signals in spatial distance decay can inform how change over time is modified by spatial relationships, something increasingly needed in an era of accelerating global change (Chen *et al*. 2025; Dallas *et al*. 2024; Kitchel *et al*. 2025; Pinsky *et al*. 2025). We detected no temporal signal in elevation distance decay across three facets of diversity using 95% credible intervals. If we expand our credible intervals to 90%, we detect a trend of biotic homogenization in species identities but not functional traits or bioclimatic niches on the highest elevation peak (Figure S1). We think this homogenization trend may be largely driven by the increase in presence and abundance of one species in particular: *Androsace septentrionalis* L.. *Androsace septentrionalis* is a diminutive short-lived annual alpine plant with a circumboreal distribution and has traits and a bioclimatic niche that match the local community on White Mountain Peak (small thin leaves, short height and a cold, wet bioclimatic niche). Increasing abundance of *Androsace septentrionalis* on lower elevation transects on White Mountain Peak has made lower transects more similar to the higher transects, homogenizing taxonomic identities but not functional traits or bioclimatic niches across the elevation gradient. This decoupling of taxonomic and functional responses illustrates how increased presence/abundance of a functionally redundant species, one whose traits and climatic niche fall within the existing range of community strategies, can alter species composition without shifting functional or bioclimatic similarity. Previous analyses of alpine plant community change in the White Mountains, CA, USA found limited plant community change on summits (Goff *et al*. 2025). We add to this understanding by showing that even below summits, where dispersal and establishment lags may be reduced due to closer proximity to sources of lower elevation species and less extreme environmental conditions, plant community compositional change was limited. More broadly, this means that there has not yet been sufficient climate change to elicit plant community change, or the time scale for plant community changes in xeric alpine areas is longer than 5-20 years due to lags in dispersal, establishment, and extinction.

In conclusion, the distance-decay analysis framework was effective in elucidating patterns of alpine plant community change across elevation gradients sampled 3-4 times between 2007 and 2024, including multiple facets of biodiversity. There are several exciting avenues for future research. First, including the rate of change of locally measured climate variables and comparing these to the rate of community compositional change would help to reveal the amount of environmental change needed to produce a given amount of community change. Second, other facets of diversity could be included in the distance decay framework, such as phylogenetic diversity (Faith 1992), the diversity of biotic interactions (Gaüzère *et al*. 2022), and rarity (Riva & Mammola 2021). Including and comparing a wider set of facets of diversity could help evaluate alternative hypotheses about community assembly and membership. Third, we did not investigate geographic signals in temporal decay because each transect only had 3-4 surveys. However, as we continue to collect data on these transects, we will be able to examine the rate of community change at the subalpine ecotone versus on summits, a pressing question in alpine ecology. As studies of distance-decay relationships accumulate at a variety of spatial and temporal scales, we will gain a better understanding of the drivers and generality of these relationships and be better equipped to apply the distance-decay analysis framework to an era of rapid biotic change.

## Acknowledgements

Brian Enquist, Malin Pinsky, Tad Dallas, Lauren Holihan, Cleber Ten Caten, and Will Petry provided inspiration and valuable feedback on this manuscript. We thank the hundreds of GLORIA Great Basin volunteers for collecting plant community data on these transects, Sylvie Amstutz for assistance in functional traits data collection, and the Plant Functional Traits 7 course for training on functional traits best practices. Aeran Coughlin, Mia Weigmann, Brooke Caldwell, Mataeus Funderburk, Sulma Correa and Emma Wilson contributed to data entry and management. Permits and research support were provided by the Inyo National Forest. Leaf material was collected under US Forest Service collection permits 24-JK-05, 23-BE-07, and 22-BE-03. This work was funded by the North Carolina State University Kenan Institute for Engineering, Technology & Science Climate Leaders Program (KAG and SNS), the Southeast Climate Adaptation Science Center Climate Change Research Fellowship (Cooperative Agreement #G23AC00548, KAG), White Mountain Research Center minigrant (KAG), Botanical Society of America Bill Dahl Graduate Student Research Award (KAG), Northern California Botanists Research Scholarship (KAG), a Mary DeDecker Botanical Grant from the California Native Plant Society Bristlecone Chapter (KAG), NCSU Faculty Research and Professional Development Grant (SNS), and the US Department of Agriculture’s National Institute of Food and Agriculture Research Capacity Fund (Hatch award number 7002993, SNS). Any use of trade, firm, or product names is for descriptive purposes only and does not imply endorsement by the U.S. Government.

