## supplemental materials for "Temporal stasis in three facets of biodiversity reveals mechanisms of community assembly across elevation gradients"

### Supplemental Methods

### Appendix S1: Supplemental Methods - Plant Functional Traits

Study Area

In 2022, we set out to investigate changes over time in arid alpine plant communities from a functional trait perspective, constructing a functional traits data set focusing on seven alpine summits in the White Mountains, CA: Sage Hen Flat (SHF), Patriarch Grove (PGS), Cottonwood South (CWS), Sheep Mountain East (SME), Mount Barcroft (BAR), Research Natural Area (RNA) and White Mountain Peak (WMT) (Bishop 2011). The White Mountains are in the rain shadow of the Sierra Nevada on the California/Nevada border, U.S.A, and receive most of their annual precipitation as winter snow and occasional summer monsoon. Summit elevations range from ~3258 m a.s.l (SHF) to ~4285 m a.s.l. (WMT). These seven summits have been studied using the GLORIA protocol (Pauli et al. 2015) every five years since 2004. We created a list of all species found in the summit area section data type of the GLORIA protocol. This is the largest and most inclusive scale, consisting of a census of all species and their qualitative abundance on all aspects within 10 m vertical distance of the highest summit point. This yielded 79 target species, including their qualitative abundance and on which peaks they were found, using the most recent GLORIA survey (2019/2021).

In 2023, a second field campaign was conducted with the goal of collecting functional trait data for all species present on GLORIA Great Basin downslope transects on Campito Mountain (CPT), Patriarch Grove (PGS), Sheep Mountain East (SME), Mount Barcroft (BAR), and White Mountain Peak (WMT). These transects extend from just below the summits to where the terrain begins to flatten and are composed of 8-14 transects that are 100m long and 1m wide. These belt transects are evenly spaced 25m vertical distance from one another on a single peak and are arranged on contour on E and SE aspects. We created a list of species on downslope transects that we had not previously sampled in 2022, which yielded 85 species. We did not set up each transect during functional trait data collection, and did our best to avoid sampling traits from the 1m wide belt transects. We aimed to collect functional traits from species over their elevational range and recorded the elevation of each individual sampled. In 2024, we returned and sampled traits from species we missed in 2023.

Our overarching aim was to measure five functional traits: maximum vegetative height, maximum generative height, leaf area, leaf dry matter content and specific leaf area for all species present in the downslope transects community data. We prioritized species with the highest cover in each year, aiming to capture species that made up at least 80% of the total cover in each summit community in each survey year (Steinbauer et al. 2022). Due to time constraints, we endeavored to create locally collected average trait means for each trait for each species. If a species occurred on multiple peaks, we attempted to collect trait data from an equal number of individuals from each peak. We aimed to sample at least 10 individuals per species across all seven peaks. For example, *Elymus elymoides* occurred on all seven peaks and were collected 1-2 times on each peak.

<u>Field Methods:</u>

Field surveys were conducted between 05 July 2022 to 25 July 2022, 02 July to 08 July 2023, and 11 July 2024 to 15 July 2024, when alpine plants in the White Mountains were at peak bloom, allowing for the best possible identification, and when plants had mostly finished vegetative growth and were allocating energy and resources to flowers and fruits. We used established protocols for functional trait measurements following Perez-Harguindeguy et al. 2016. We measured maximum vegetative and generative height on reproductive individuals in the field using a ruler, measuring between the ground and the tallest vegetative or reproductive structure, respectively. We did not stretch, pull or manipulate the plant when collecting these data, seeking to capture how far the plant projected into the sky had we not been there measuring it. After measuring these heights, we carefully collected 1-4 mature, fully-grown sun leaves that did not show signs of herbivory or pathogen damage from the same plants on which we measured heights. For all leaf collections and measurements, we included petioles and stipules where present. For grasses, we used fully formed blades from the ligule to the tip of the blade. Leaves were immediately placed in a moist paper towel in a soft-sided cooler with an ice pack to avoid dehydrating before they could be processed. We spent less than 6 hours per day in the field, generally collecting during the morning hours, in order to keep leaves in their fully hydrated state. The collection number and the height measurements in centimeters were recorded on a data sheet, in order to associate each individual's trait measurements throughout the study.

Leaf samples were processed at the lab at Crooked Creek Field Station, which is a maximum of one hour walk or drive from sampling sites. We used two SmartWeigh GEM20 portable balances to measure wet leaf weight, to the nearest 0.001 g. Leaves were then scanned using two Canon CanoScan LIDE 300/400 scanners, which were connected to laptops. After weighing and scanning, leaves were placed into coin envelopes and 25 envelopes at a time were placed into a ziplock bag containing ~500 g of silica gel. Samples remained in silica gel until they were unpacked and dried in a drying oven.

Laboratory Methods:

On 23 August 2022, 05 September 2023 and 01 September 2024, leaf samples were removed from bags of silica, organized alphabetically by species, and placed upright in paper bags. This took place over the course of 3 hours on a single day in the lab, to avoid leaves rehydrating by sitting on the lab bench at ~35% RH. Leaves were initially dried for 72 hours at 60 °C using a Fisher IsoTemp Drying Oven. Samples were removed from the oven and weighed periodically, to check for changes in moisture content. Leaf samples were stored in the drying oven during the weighing process. Dry leaf weight was measured to the nearest 0.0001 g using Mettler Toledo XSR104 microbalance. We counted the number of leaves in the envelopes before weighing to check if any of the very small leaves had been lost in transport.

Data Processing and Analysis:

We cropped images in R using the functions “image_crop” and “image_write” from the R package *magick* (Ooms 2025), in a custom function. This function pulled scans from one folder, cropped the plant scan out of the taped area, and wrote the new image to another folder. We used ImageJ (Schneider et al. 2012) to measure the distance from the upper edge, distance from the left edge, width and height of the scans in order to specify the crop area. We manually checked each crop to make sure no tape was left in the scans following this process. Next, we used ImageJ and the function “run.ij”, part of the *LeafArea* package in R (Katabuchi 2015) to automate the leaf area measurements. We manually measured a known area on a ruler in the scan using ImageJ, and calibrated the known distance to a number of pixels (5 cm = 592 pixels in our scans). Due to the very small size of some of our leaves, we could only filter out objects smaller than 0.0001 cm^2 using the *low.size* argument in the “run.ij” function. We saved the threshold images that ImageJ uses to measure leaf area, so that we could manually check the leaf area measurements for potential problems.

Several species of alpine plants have silvery long-hairy to pubescent leaves, which is challenging to measure the area of using threshold methods because the contrast between the leaf and the scanner platen is low. We manually checked all threshold images from imageJ, and where pieces of leaves or petiole were missing, we manually colored in the missing parts of the leaf using an Ipad Pro 10" and Apple Pencil. To do this, we used the eyedropper tool to use a color that was similar to that found in the rest of the leaf that was picked up by the automated leaf area methods. These manually colored leaves had their leaf areas re-measured using the methods in the previous paragraph, and threshold images were checked again to make sure the manual coloring was successful.

Trait values for each collection, and each species, were calculated in R. Because of the size of our leaves and sensitivity of our field instruments, we often measured several leaves of the same individual. For this reason, we noted the number of leaves during the wet mass and leaf area recording, and the number of leaves during the dry mass recording. In a few rare cases, these numbers did not match, likely due to very small leaves falling out of the coin envelopes. To account for this, we calculated the leaf wet mass for a single leaf by dividing the number of wet leaves by the leaf wet mass. We repeated this process for the leaf dry mass (dividing by the number of dry leaves), and the leaf area (dividing by the number of wet leaves). Next, we calculated leaf mass area (LMA - $g/cm^2$; dry leaf mass divided by leaf area), specific leaf area (SLA - cm/g; leaf area divided by dry leaf mass) and leaf dry matter content (LMDC - g/g; leaf dry mass divided by leaf wet mass) for each individual.

**Appendix S2: Supplemental Methods - Occurrence Data and Bioclimatic Niche**

To understand how communities vary across elevation and temporal gradients in terms of their bioclimatic niches, we constructed climatic niche means for each species. We began the process of constructing climatic niche means by assembling occurrence data from each species represented in our community dataset. Occurrence data for each species was downloaded from the Global Biodiversity Information Facility (Derived Dataset GBIF.org Filtered Export of GBIF Occurrence Data., 2024) using the package "rgbif" in R (Chamberlain et al., 2024). In total, we downloaded 200,778 occurrences for 146 species found in our dataset on plant communities in the White Mountains, CA. We only downloaded occurrence data that had latitudinal and longitudinal coordinates and voucher species found in herbaria. We excluded any occurrence data that had known geospatial issues according to GBIF, did not have an herbarium record, or was collected before 1948. We filtered the data download to 15 of 224 total data fields, including: species name, latitude and longitude, country code, individual count, GBIF ID, family, taxon rank, coordinate uncertainty in meters, year, basis of record, institution code, dataset name, state or province, and accepted scientific name.

We began cleaning occurrence data using automated methods contained in the R Package “CoordinateCleaner” (Version 3.0.1, Zizka et al., 2019). Specifically, we used the CoordinateCleaner tests “capitals”, “gbif”, “institutions”, “urban”, and “seas” to identify and flag points that fell in capital cities, the GBIF Headquarters in Copenhagen, Denmark, biodiversity institutions like museums, urban areas, and oceans, respectively. Additionally, we used the CoordinateCleaner package tests “equal” and “zeros” to identify and flag points that contained incorrectly entered coordinates, such as latitudes and longitudes that were equal to each other or entered as zero. These seven CoordinateCleaner tests flagged and excluded 5,510 observations (2.74% of the initial 200,778 observations), and the remaining dataframe contained 195,269 observations. Next, we removed occurrences with coordinate uncertainty over 1 kilometer to align the data with the resolution of the climate data. A total of 160,695 occurrences remained after this step. Data with imprecise coordinates (zero decimal places) or data lacking GBIF IDs were omitted. A GBIF ID is the unique number associated with a GBIF occurrence record. It is important that every occurrence has a GBIF ID in order to locate an occurrence on the website and verify its accuracy. Finally, we filtered all occurrences after the year 2000 to attempt to ensure that they were collected with a GPS unit. This left us with 62,017 occurrences. All species used in analysis had at least three occurrences.

Following our automated cleaning steps, we created maps of the occurrences for each species (Deckmyn et al. 2024). We used these maps for manual cleaning. Occurrences that fell far outside of a given species’ typical range were flagged for further inspection. During the manual cleaning process, we isolated the points by filtering based on approximate latitude and longitude. Then, we entered the GBIF IDs from those points into the GBIF website, checking for an accurate herbarium specimen, consistency with Flora of North America and iNaturalist species distributions, or coordinates outside of North America. Occurrences that we could not confirm were removed from the final occurrence dataset.

After saving the final cleaned occurrence dataset, we needed to overlay it with climate data in order to define the average climatic conditions of each species' geographic distribution. We downloaded all 19 available global bioclimatic variables from 1981 to 2010 using Version 2.1 of CHELSA Climate (Brun et al., 2024). Each bioclimatic variable was downloaded as an individual TIF file, a raster layer at a resolution of 30 arc sec. These rasters were then stacked and cropped to the approximate geographic extent of North America, using the R package "terra" (Version 1.8-42, Hijmans et al. 2025). Importantly, we converted the coordinate reference system (CRS) of the new spatial occurrence points to WGS 1984 so that it matched the map projection used in the CHELSA Climate data. We also converted the occurrences into spatial points using the st_as_sf() function within the sf package, setting the latitude and longitude columns from the occurrence data to be the coordinates associated with each point. This allowed us to plot the occurrences as spatial points on top of the climate rasters and extract the climate values to each occurrence based on the matching coordinates. The end product was a large SpatVector with climate values assigned to each species occurrence. We thinned the occurrences to one occurrence per species per grid cell in order to reduce duplication and sampling bias. Finally, we calculated niche means for five bioclimatic variables for each species: mean annual temperature (°C), mean annual precipitation (mm), isothermality (mean of monthly max - min), temperature seasonality (°C), and precipitation seasonality (mm).

### Appendix S3: Supplemental Methods - Taxonomic Harmonization

In the community dataset, the abundance of dummy species was the sum of all species aggregated to create that dummy species, capped at a maximum of 10 out of 10 segments. For functional traits and bioclimatic niches, we averaged functional traits and bioclimatic niche means. For example, we created the dummy species "Antennaria sp." from all species of *Antennaria* Gaertn. recorded across all peaks and years (4 species) by summing the abundance in the community data, and averaging the trait and bioclimatic niche data across the original four species. We repeated this process for 4 species of *Boechera* Á.Löve & D.Löve and four species of *Draba* Dill. ex L.. Finally, we aggregated *Poa secunda* Zea ex Roem. & Schult. and *Poa glauca* Vahl into the dummy species "Poa secunda/glauca", *Poa keckii* Soreng and *Poa lettermanii* Vasey into "Poa keckii/lettermanii", *Castilleja nana* Eastw. and *Castilleja pilosa* (S.Watson) Rydb. into "Castilleja nana/pilosa", and *Festuca minutiflora* Rydb. and *Festuca brachyphylla* Schult. & Schult.f. into "Festuca minutiflora/brachyphylla".

### Supplemental Figures

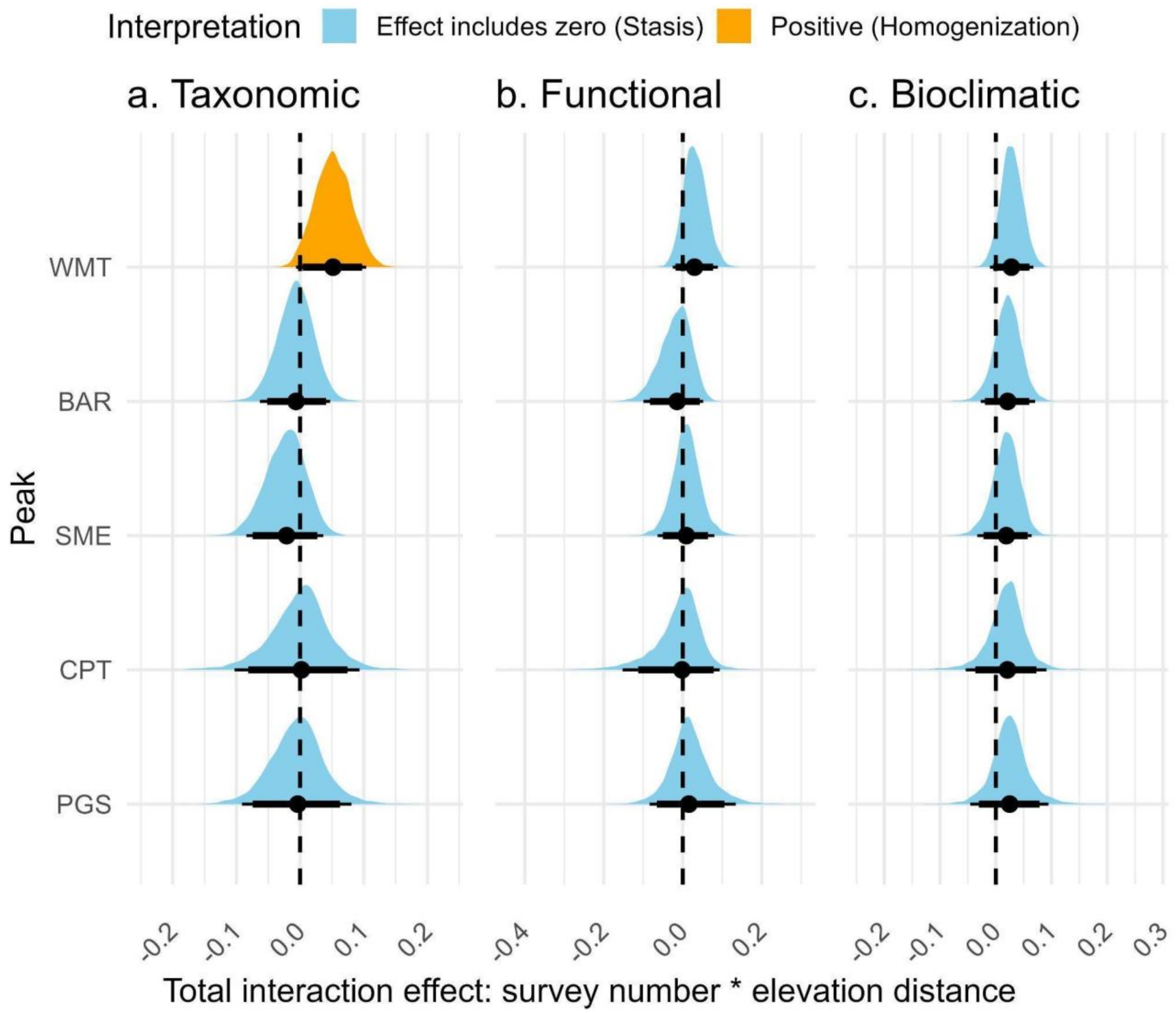


**Figure S1:** Posterior probability distributions for the total interaction effect of elevation distance and survey number, computed via the sum of the elevation distance * survey number fixed effect and each peak-specific interaction effect, across three facets of diversity. On peaks where the total interaction effect 95% credible interval overlaps zero, we find evidence for no change in the elevation distance decay-relationship, or community stasis. When the 90% credible interval is used, we find evidence for biotic homogenization of taxonomic identities on the highest elevation peak, White Mountain Peak. 90% and 95% credible intervals are shown below each distribution.

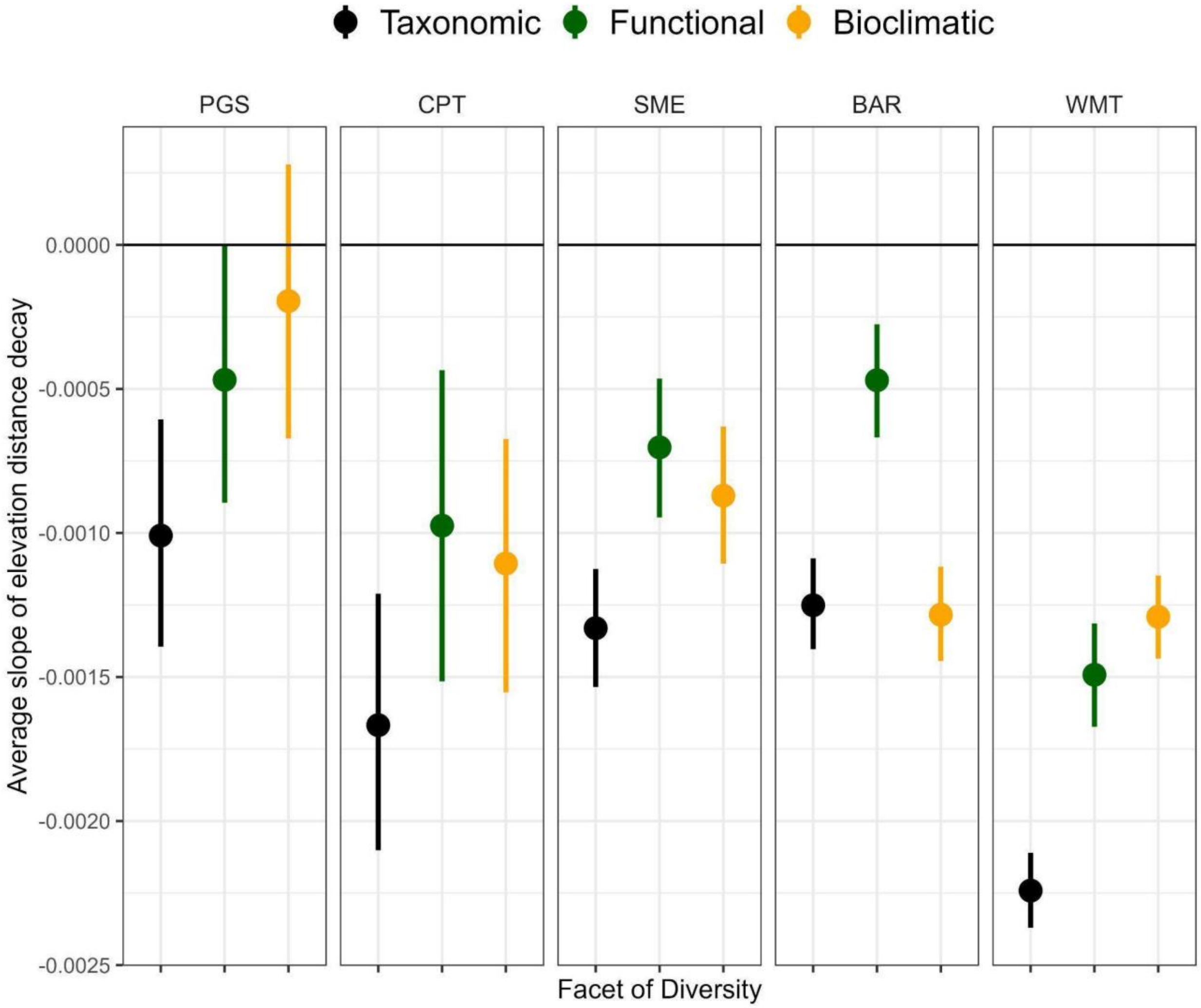


**Figure S2:** Slopes of elevation distance-decay relationships across three facets of diversity: taxonomic (species identities), functional (functional diversity in vegetative height, SLA and LDMC) and bioclimatic niche (suite of climate variables representing averages, variability and extremes), holding survey year at a fixed average value. Line ranges show 95% credible intervals. Facets are arranged by median peak elevation, with the lowest peak on the left and the highest peak on the right.

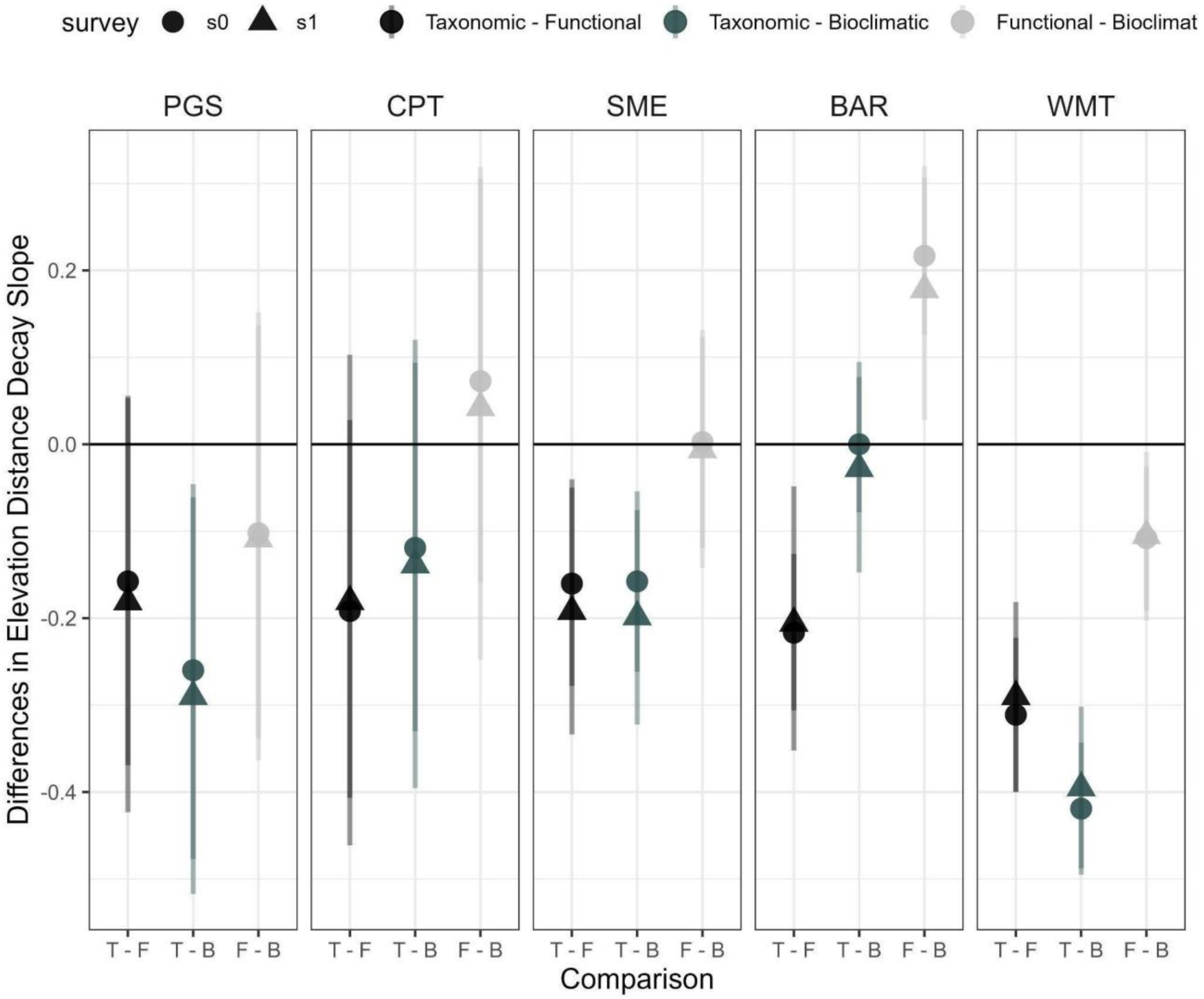


**Figure S3:** Posterior differences in elevation distance-decay slopes between biodiversity facets (colors), including the effect of time (“survey”, shapes). Points show posterior means; lines show 95% credible intervals. Positive values indicate shallower slopes for the first facet; negative values indicate steeper slopes. 95% credible intervals for a baseline survey and a one-unit increase in survey number overlap, and thus survey number does not modify the slope differences among facets.
